# Impact of temperature and photoperiod on survival and biomarkers of senescence in common woodlouse

**DOI:** 10.1101/433011

**Authors:** Charlotte Depeux, Ascel Samba-Louaka, Christine Braquart-Varnier, Jérôme Moreau, Jean-François Lemaître, Tiffany Laverre, Hélène Pauhlac, François-Xavier Dechaume-Moncharmont, Jean-Michel Gaillard, Sophie Beltran-Bech

## Abstract

Most living organisms display a decline in physiological performances when ageing, a process called senescence that is most often associated with increased mortality risk. Previous researches have shown that both the timing and the intensity of senescence vary a lot within and among species, but the role of environmental factors in this variation is still poorly understood. To fill this knowledge gap, we investigated the impact of environmental conditions on the strength of senescence using an experimental design applied to a population of common woodlouse *Armadillidium vulgare* intensively monitored in the lab. Cellular senescence biomarkers are available in woodlouse and are age-related. These biomarkers provide relevant biomarkers to test the impact of environmental conditions, through changes in temperature and photoperiod, on individuals of the same age maintained in different environmental conditions. We found different effects of the environmental changing: the increasing of day light modification leaded the same effect as age on our senescence biomarkers while temperature modifications leaded the opposite effect as age on the β-galactosidase activity and cell size. We also demonstrated the existence of sex-specific responses to changes in environmental conditions. By using an experimental approach and biomarkers of senescence in woodlouse, we show that environmental conditions and sex both shape the diversity observed in senescence patterns of woodlouse and underline the importance of identifying senescence biomarkers to understand how environmental conditions influence the evolution of senescence.

## 1. Introduction

Senescence is generally defined as a progressive decline in physiological performances that leads to a decrease in the probability to reproduce (i.e. reproductive senescence) or survive (i.e. actuarial senescence) with increasing age (Monaghan et al., 2008). This process is nearly ubiquitous in the living world ((Nussey et al., 2013)) but displays a tremendous diversity of patterns across the tree of life (Jones et al., 2014; Shefferson et al., 2017). Whatever the studied trait, both timing and intensity of senescence strongly vary across species (Jones et al., 2014; Nussey et al., 2011), populations (Hassall et al., 2017; Tidière et al., 2016), and individuals (Bérubé et al., 1999). Many studies aiming to understand the diversity of senescence patterns at different levels of the biological organization have suggested that environmental conditions are likely to play a significant role (Fontana et al., 2010; Martin et al., 1996). For instance, resource competition in early life can strengthen both actuarial and body mass senescence in wild populations of mammals (Nussey et al., 2007; Beirne et al., 2015). However, environmental conditions can potentially influence senescence in a sex-specific way, as evidenced for other life-history traits. For instance, in the neriid fly (*Telostylinus angusticollis*), a dietary restriction caused the complete female infertility, whereas in males, the negative effect of dietary restriction on reproduction was effective only when they received a rich larval diet and when they were housed with females (Adler et al., 2013). In the Alpine marmot, (*Marmota marmota*), the social environment lead strongly different actuarial senescence patterns between males and females (Berger et al., 2018). Sex-specific effects of environmental conditions on senescence thus need to be investigated. Moreover, to thoroughly understand how environmental conditions modulate observed patterns of senescence, their specific impact on organisms must be separately quantified. Here, we provide such a study by investigating cellular senescence in the common woodlouse *Armadillidium vulgare*. This terrestrial isopod can live up to three years (Paris and Pitelka, 1962) and is highly sensitive to environmental conditions, especially when they involve changes in photoperiod and temperature. In fact, these parameters are closely related to reproduction and water loss (Brody et al., 1983; Hassall et al., 2018; Mocquard et al., 1989, 1980; Smigel and Gibbs, 2008; Souty-Grosset et al., 1988). Overall, woodlouse can be easily controlled and monitored in the laboratory and thereby constitutes a very relevant model to test whether and how environmental conditions impact senescence patterns.

Biomarkers of senescence correspond to biological parameters that allow predicting the functional capability of an organism better than its chronological age (Baker and Sprott, 1988). At the cellular scale, senescence corresponds to the cellular deterioration leading stop of the cellular cycle (Campisi and di Fagagna, 2007). As ageing is associated with cellular senescence (Herbig et al., 2006; Lawless et al., 2010; Wang et al., 2009), cellular biomarkers provide reliable metrics to study senescence. One of the most popular biomarker of senescence is based on the lysosomal activity of the β-galactosidase enzyme, which increases when the cell enters in senescence (Dimri et al., 1995; Itahana et al., 2007). The activity of the β-galactosidase has mostly been used to study the senescence of mammalian cells (Gary and Kindell, 2005), but has also been successfully used to detect both senescence in honeybees (Hsieh and Hsu, 2011) and effect of lower temperature on senescence of the short-lived fish *Nothobranchius furzeri* (Valenzano et al., 2006). Likewise, the decline in immune performance with increasing age (i.e. immunosenescence) can also provide a suitable biomarker of senescence. A diminution of the number of effective immune cells has thus been reported in wild vertebrates (Cheynel et al., 2017) but also in invertebrates including mosquitoes *Aedes aegypti* (Hillyer et al., 2004) and crickets *Gryllus assimilis* (Park et al., 2011). In this later species, the immunosenescence also involves a decrease of the melanotic module formation, which increases damage of immune cells, and then modifies the immune cell composition (Park et al., 2011). Moreover, senescent cells are bigger than non-senescent cells (Hayflick, 1965; Rodier and Campisi, 2011). Overall the viability and the size of immune cells reliably indicate the level of individual immunosenescence. In biomedical research, biomarkers of immunosenescence are routinely used to assess the role of stressful environmental conditions on ageing (Piazza et al., 2010).

In this study, we used measurements of senescence biomarkers specific to *A. vulgare* (Depeux et al., 2019) as metrics to quantify the impact of changes in temperature and photoperiod on individuals of the same age experimentally maintained in different conditions of temperature or photoperiod. We expected the effect of temperature and photoperiod to shape specific variations on the senescence biomarkers and survival in female and male woodlice.

## 2. Materials & Methods

### 2.1 Biological model

Individual *A. vulgare* used in the following experiments were derived from wild populations and have been monitored in lab conditions over ten years. Individuals have been maintained on moistened soil with the natural photoperiod of Poitiers (France 46.58°N, 0.34°E, 20°C) at 20°C with food (i.e. dried linden leaves and carrots) *ad libitum*. Crosses were monitored to control genetic diversity. For each clutch obtained, individuals were sexed, and brothers and sisters were separated to ensure virginity.

In common woodlouse, individuals moult throughout their lives according to a molting cycle (Lawlor, 1976). At 20°C, they approximately moult once per month (Steel, 1980) and all the cells of the concerned tissues are renewed. However, the brain, the nerve cord, part of the digestive tract and gonads are not part of tissues renewed during molting and are therefore good candidates for tissue-specific study of senescence in this species. In addition to molting regularly, the female woodlouse exhibits specific molts related to reproduction (Moreau and Rigaud, 2002). These molts and more generally the onset of reproduction are influenced by environmental conditions: both increased temperature and longer days stimulate the onset of reproduction (Mocquard et al., 1989). The reproductive period occurs during spring.

### 2.2 Experimental design

We tested the effect of environmental factors on senescence in *A.vulgare* on the survival and on our senescence biomarkers (Figure 1). The different protocols were applied to males and females separately to assess the effect of sex on senescence patterns.

**Figure 1.**
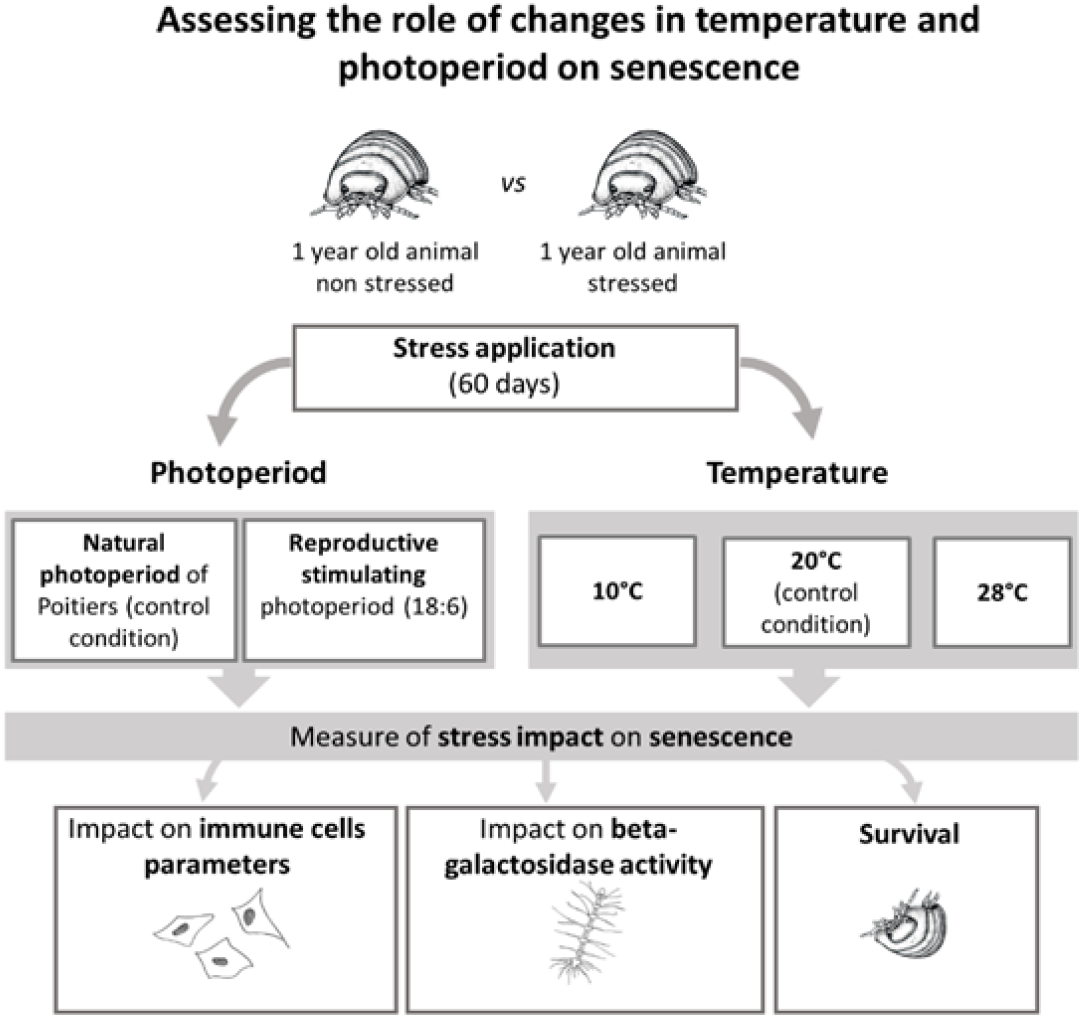
Experimental Design. 600 individuals were used to test the impact of changes in temperature and photoperiod on survival and biomarkers of senescence: 120 individuals by experimental (or control) condition composed of 60 females and 60 males. We estimated 60 days after exposure to different temperature and photoperiod conditions the survival of these 120 individuals (by condition). More specifically, 30 (15 males and 15 females) individuals were sampled to measure the impact of changes in temperature and in photoperiod on immune cells and on β-galactosidase activity using the photoperiod experimental or control protocols and the temperature protocols at 10°C and 20°C. For the temperature protocol at 28°C, fewer females (25) and males (20) were sampled because of the lower survival rate under these conditions.

To measure the effect of temperature and photoperiod on senescence, we used 120 individuals by experimental (or control) condition (i.e. natural photoperiod, stimulating photoperiod, 10°C, 20°C and 28°C) composed of 1 year old individuals (60 females and 60 males). At the beginning of the experience, the same quantity of soil and 5g of dry food were weighed, rehydrated, and disposed in boxes (length x width x height: 26.5 × 13.5 × 7.5 cm). To test the impact of photoperiod, we exposed *A. vulgare* to either natural photoperiod (corresponding to the photoperiod observed from January to March at Poitiers) as controlled conditions or a photoperiod stimulating woodlouse reproduction (18:6 D/N) as experimental conditions at a temperature of 20°C. To test the effect of temperature, we exposed *A. vulgare* to three different temperatures: 20°C corresponding to controlled conditions, 10°C corresponding to supposed stressful cold conditions, and 28°C corresponding to supposed stressful “hot” conditions. After 60 days of exposure at the five different conditions, individuals were enumerated to estimate survival. Survivors were then maintained in normal laboratory conditions (i.e. at 20°C under natural photoperiod) during two months until the measures of cellular senescence using senescence biomarkers previously developed. The survival at different temperatures was estimated from 120 individuals. To estimate the cellular senescence on biomarkers of senescence, 30 individuals from each sex were collected from each environmental condition except at 28°C, when high mortality forced us to collect only 20 males and 25 females. The measure of immune cell parameters was realized at the individual scale when the measurement of the β-galactosidase required a pool of 5 nerve cords (i.e. from 5 individuals) to obtain enough biological material. In this way, the hemolymph sampling was achieved in each individual before dissection of the animals and reunification of the nerve cords of 5 animals.

### 2.3 Measure of immune cell parameters

To study the impact of environmental conditions on the immune parameters, 3 μL of haemolymph were collected per individual. A hole was bored in the middle of the 6^th^ segment and 3 μL of haemolymph were collected with an eyedropper and deposited promptly in 15 μL of MAS-EDTA (EDTA 9 mM, Trisodium citrate 27 mM, NaCl 336 mM, Glucose 115 mM, pH 7, (Rodriguez et al., 1995)). Then, 6 μL of Trypan blue at 0.4% (Invitrogen) were added to permit the coloration of dead cells. Thereafter, 10 μL of this solution were deposed in (Invitrogen Coutness®) counting slide (Thermofisher). The immune cell density, the immune cell viability and the immune cell size were evaluated using an automated Cell Counter (Invitrogen Countess®).

### 2.4 Measure of β-galactosidase activity

To test the impact of environmental conditions on β-galactosidase activity, individuals were dissected separately in Ringer (Sodium Chloride 394 mM, Potassium Chloride 2 mM, Calcium Chloride 2 mM, Sodium Bicarbonate 2 mM) and nerve cord was removed. Nerve cords were chosen because they are not regenerated during molting. To obtain a sufficient quantity of protein, we made pools of five nerve cords (from five different individuals of the same age). The five nerve cords were filed in 500 μL of Lyse Buffer 1X (CHAPS 5 mM, Citric acid 40 mM, Sodium Phosphate 40 mM, Benzamidine 0.5 mM, PMSF 0.25 mM, pH = 6) (Gary and Kindell, 2005). Samples were centrifuged at 15000g at 4°C for 30 minutes. The supernatant was taken and kept at −80°C until its utilization. The protein concentration was determined by the BCA assay (Thermofisher) and were homogenized at 0.1 mg/mL.

The β-galactosidase activity was measured as previously described by Gary and Kindell (2005). Briefly, 100 μL of protein extract at the concentration of 0.1 mg/mL were added to 100 μL of reactive 4-methylumbelliferyl-D-galactopyranoside (MUG) solution in a 96 well-microplate. The MUG reactive, in contact to β-galactosidase, leads by hydrolysis to the synthesis of 4-methylumbelliferone (4-MU), which is detectable using fluorescent measurement. Measures were performed by the multimode microplate reader Mithras LB940 HTS III, Berthold; excitation filter: 120 nm, emission filter 460 nm, for 120 minutes. Two technical replicates were measured for each pool.

### 2.5 Statistical analysis

The β-galactosidase activity was analyzed with linear mixed effect models using the R package lme4 (Bates et al., 2014). As two technical replicates were measured for each pool, the model including the pools fitted as a random effect and age, photoperiod or temperature, and sex and their two-way interactions as fixed factors. Linear models with Gaussian distribution were fitted to analyze variation in the cell size and viability. For the cell density, a linear model of the cell number (log-transformed, (Ives and Freckleton Robert, 2015) was fitted. Survival in different conditions of temperature and photoperiod was analyzed with generalized models (with a binomial error). All results were obtained by stepwise backward selection model. The effect size of temperature and photoperiod on the β-galactosidase activity and immune parameters was measured by rescaling and standardized slopes were reported as a measure of effect size (Schielzeth, 2010).

## 3. Results

### 3.1 Photoperiod

Stimulating photoperiod did not have any detectable influence on survival in each sex (X^2^_1_= 0.20, p=0.65 and X^2^_1_=1.96, p=0.16 for males and females, respectively) but led to a higher β-galactosidase activity, (X^2^_1_=3.86, p=0.05) and did not influence the size (F_1,108_=0.264, p=0.61) density (F_1,108_=0.54, p=0.54) or viability (F_1,108_=0.83, p=0.36) of immune cells (Figure 2).

Sex did not impact significantly the β-galactosidase activity (X^2^_1_=1.96, p=0.16), the size (F_2,108_=0.32, p=0.57), density (F_2,108_=0.41, p=0.52) or viability (F_2,108_=1.50, p=0.22) of immune cells (Figure 2).

**Figure 2.**
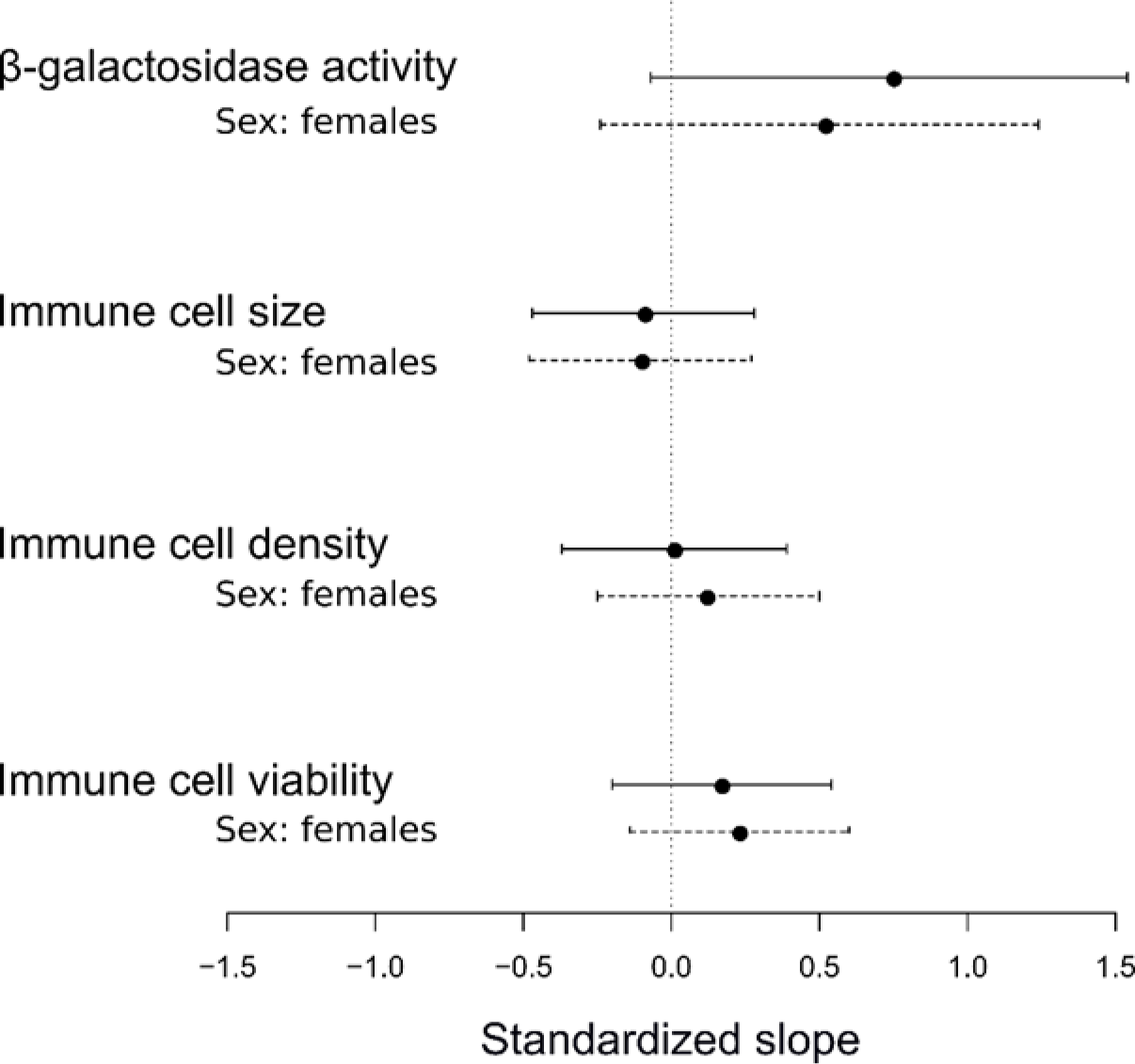
Effect size of photoperiod stress on the β-galactosidase activity, immune cell size, immune cell density and immune cell viability. This figure synthetizes the effect size with bootstrapped 95% CI of non-optimal photoperiod (solid lines), with the sex effect (dotted line) for each senescence biomarkers studied. In each case, the effect is relative to the control condition (natural photoperiod for stimulating photoperiod) and for the sex, females were compared to males.

### 3.2 Temperature

At 28°C, the survival of animals decreased (X^2^_1_=47.48, p<0.001). However, at 10°C, survival of animals was not impacted (X^2^_1_=0.42, p=0.51). Temperature had an influence (Figure 3) on the β-galactosidase activity (X^2^_2_=12.2, p=0.002), on the cell size (F_2,291_=7.96, p=0.01), but also on cell density (F_3,290_=9.24, p=0.001) and viability (F_3,290_=19.37, p<0.001). In fact, β-galactosidase activity showed lower values in the two stressful temperature conditions (i.e. 10°C (X^2^_1_=9.67, p=0.002) and 28°C (X^2^_1_=3.85, p=0.05)) than in the control condition (i.e. 20°C). Under stressful temperature of 10°C cells were smaller than in temperature 20°C (F_2,207_=9.24, p=0.006). The density of immune cells was lower under the stressful temperature 10°C (X^2^_1_=18.96, p<0.001) and 28°C (X^2^_1_=7.06, p=0.008) as the cell viability (Respectively: X^2^_1_=15.98, p<0.001; X^2^_1_=15.80, p<0.001).

A sex effect was also detected with a higher β-galactosidase activity in females than in males (X^2^_1_=16.95, p<0.001, Figure 3). The cell density was higher in females (F_3,290_=4.01, p=0.05) as was the cell viability (F_3,290_=19.61, p<0.001) (Figure 3). No detectable difference occurred between females and males in cell size (F_3,290_=0.46, p=0.49, Figure 3).

**Figure 3:**
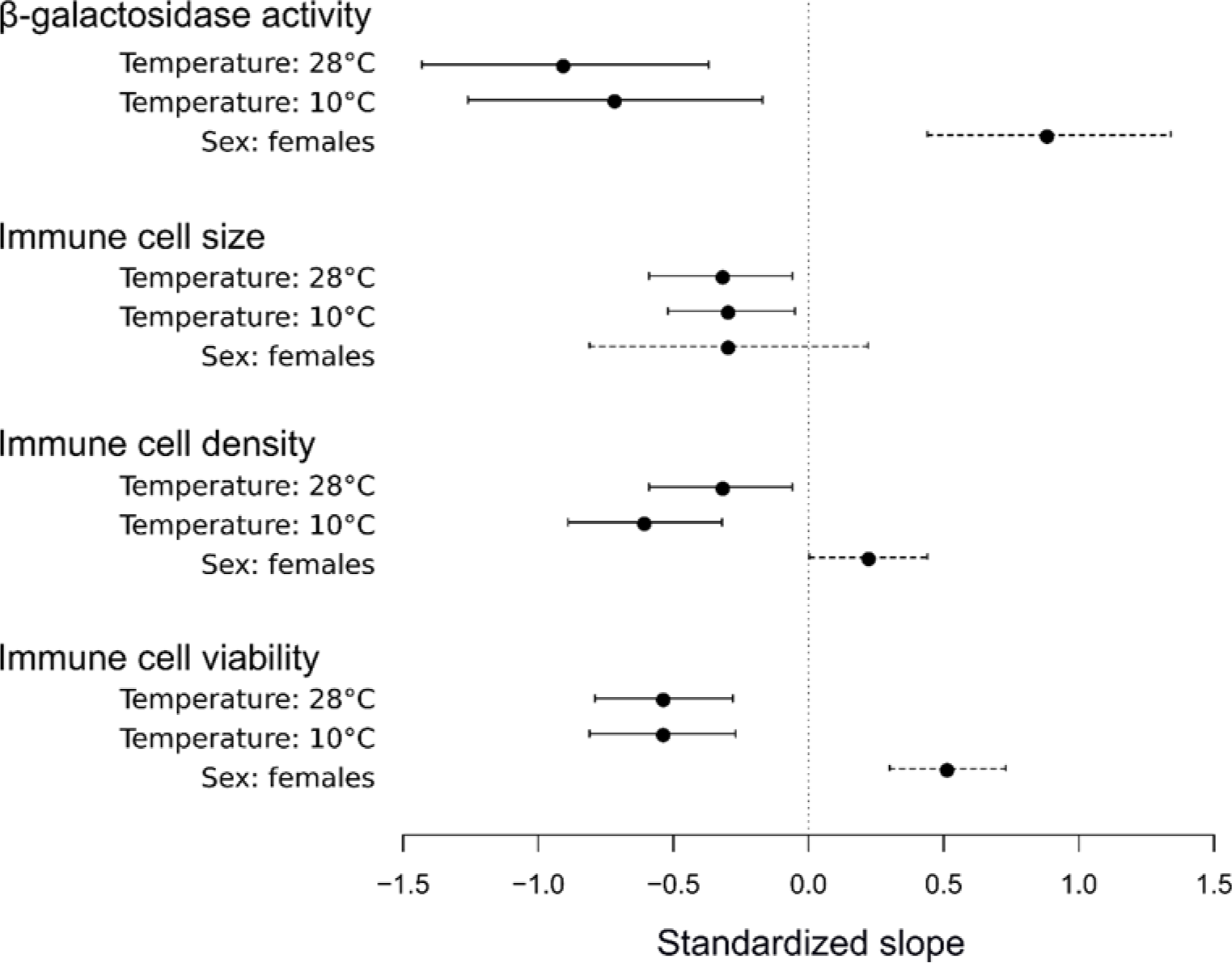
Effect size of temperature stresses on the β-galactosidase activity, immune cell size, immune cell density and immune cell viability. This figure synthetizes the effect size with bootstrapped 95% CI of temperatures (solid lines), with the sex effect (dotted line) for each senescence biomarkers studied. In each case, the effect is relative to the control condition (temperature 20°C for temperatures 10°C and 20°C) and for the sex, females were compared to males.

## 4. Discussion

To understand the diversity of senescence patterns described across the tree of life, the role of interplaying environmental factors must be identified. Our experiments provide clear evidence that changes in photoperiod and temperature influence biomarkers of senescence in *A. vulgare* and temperature influences survival too. These results suggest that environmental condition likely shape senescence patterns in *A. vulgare*.

We aimed to test the impact of environmental conditions on woodlouse senescence by using two distinct environmental stressors: the photoperiod and the temperature. While the elongation of the day light caused an increase of the β-galactosidase activity without any negative impact on survival and immune cells, cold condition led to smaller cell size. At 28°C (hot condition), survival being affected, we cannot totally exclude the hypothesis that the dead individuals did not have exactly the same profile as the survivors on the measured biomarkers. Anyhow, lower and higher temperatures than 20°C seems to induce lower β-galactosidase activity and decreased cell density and viability. Results obtained in biomarkers were highly different and underlined the important role of environmental factors on senescence. In *Drosophila melanogaster*, exposition to high temperature lead the shortening of the lifespan (Garcia et al., 2010) while caloric restriction often leads to extended lifespan in different species, from rats to worms (Koubova, 2003). In our study on *A. vulgare*, we observed that different environmental stresses could also lead opposite effects on the same biomarkers. These results suggest that stress effects observed in lifespan need to be study at the cellular scale to understand phenomenon implied.

Moreover, old females had a higher β-galactosidase activity and a better immune cell density and viability than males in stressed conditions. These results could be explained by a more effective immune system in females as often observed in the living world (Nunn et al., 2009). However, previous study in *A. vulgare* suggests the opposite: in Sicard et al. (2011), one-year-old males showed a higher cell density than females of the same age. We supposed that, according to their life history traits, different gender strategies exist in *A. vulgare* as suggested previously in (Paris and Pitelka, 1962) and our study indicates that these strategies are shaped by environmental conditions.

## 5. Conclusion

Our study confirms that *A. vulgare* presents all characteristics required to study senescence. Our findings in this species support the hypothesis that the diversity of senescence patterns observed among species results from complex interactions between sex and environmental conditions. The development of a variety of biomarkers including telomerase activity and telomere size could allow getting a deeper understanding of the impact of environmental conditions on senescence patterns. Assessing how different environmental stressors influence biomarkers should help to identify the drivers of the great diversity of senescence patterns in the living world.

## Acknowledgements

We would like to thank Sylvine Durand, Isabelle Giraud and Bouziane Moumen for our constructive discussions as well as Maryline Raimond and Alexandra Lafitte for technical assistance. We would also like to thank Richard Cordaux and Xavier Bonnet for their constructive comments.

